# Epigenetic reprogramming of DCCs into dormancy suppresses metastasis *via* restored TGFβ–SMAD4 signaling

**DOI:** 10.1101/2021.08.01.454684

**Authors:** Deepak K. Singh, Eduardo Farias, Saul Carcamo, Dan Hasson, Dan Sun, Julie Cheung, Ana Rita Nobre, Nupura Kale, Maria Soledad Sosa, Emily Bernstein, Julio A. Aguirre-Ghiso

## Abstract

Disseminated cancer cells (DCCs) identified in secondary organs, sometimes before the primary tumor becomes detectable and treated, can remain dormant for years to decades before manifesting. Microenvironmental and epigenetic mechanisms may control the onset and escape from dormancy, and here we reveal how a combination of the DNA methylation inhibitor 5-azacytidine (AZA) and retinoic acid receptor ligands all-trans retinoic acid (atRA), orchestrate a novel program of stable dormancy. Treatment of HNSCC tumor cells with AZA+atRA induced a SMAD2/3/4 dependent regulation of downstream transcriptional program that restored the anti-proliferative function of TGFβ signaling. Significantly, AZA+atRA or AZA+AM80, an RARα specific agonist, strongly suppresses lung metastasis formation. The metastatic suppression occurs *via* the induction and maintenance of phenotypically homogenous dormant SMAD4+/NR2F1+ non-proliferative DCCs. These findings suggest that strategies that maintain or induce dormancy programs may be a viable alternative strategy to improve patient outcomes by preventing or significantly delaying metastasis development.

## INTRODUCTION

Metastasis is a multi-step process and is the primary cause of cancer-related deaths^1, 2^. The progression of metastasis is highly complex and depends on the cancer cells (“seeds”) and the microenvironment (“soil”) of secondary organs^3, 4^. Until recently, it was believed that cancer cells disseminated from advanced and clinically aggressive tumor lesions. However, several studies have shown that disseminated cancer cells (DCCs) can be present in secondary organs even before the primary tumor can be detected clinically^5, 6, 7, 8^. Importantly, genetically heterogeneous DCC populations appear to remain dormant for long periods (sometimes decades) and have the potential to reactivate in response to additional genetic, epigenetic, and microenvironmental changes^9, 10, 11^. These findings suggest that strategies that manage post-extravasation events, and thus DCC biology may be one of the shortest paths to changing patient outcomes by preventing or significantly delaying metastasis development.

The available experimental and human DCC data has revealed that cancer cells appear to spend long periods in a quiescent state during dormancy, which is evident by the upregulation of cyclin-dependent kinase inhibitors, like p21 and p27^9, 10, 11^. However, many studies have revealed a higher level in the hierarchy of signaling mechanisms that control dormancy where specific cues such as retinoic acid, BMP4/7, LIF, GAS6, and TGFβ2 can activate specific transcription factors (TFs) (e.g., NR2F1, RARβ, DEC2, NDRG1) that commit cancer cells to a long-lived cell cycle arrest accompanied with cell plasticity or “stem-like” programs (e.g., SOX9, SOX2)^9, 10, 11^. The signaling and transcriptional programs mentioned above involve genes controlling proliferation/quiescence, stress tolerance, survival, and pluripotency/self-renewal^12, 13, 14^. In addition, dormant cells have been shown to display a repressive chromatin state that seems long-lived, dependent on microenvironmental cues such as retinoic acid, but also reversible^14^. Thus, dormant DCCs appear to undergo a transitional epigenetic reprogramming driven by niche-derived cues.

Previously, we demonstrated the reprogrammable nature of dormancy and showed that the retinoic acid regulated nuclear receptor NR2F1 was responsible for inducing and maintaining DCC dormancy. We further showed that 5-Azacytidine (AZA) in combination with *all-trans* Retinoic Acid (atRA) could reprogram tumor cells and led to the induction of quiescence and suppressed head and neck squamous cell carcinoma (HNSCC) primary tumor PDX (patient-derived xenograft) growth *in vivo* in part *via* NR2F1 function^14^. atRA is the biologically active form of vitamin A, and it mediates its action through RARs and RXRs heterodimers^15^. It controls quiescence and differentiation during development and tissue homeostasis. For example, hematopoietic stem cells respond to atRA by entering a cell-cycle arrest and maintaining a dormant state^16^. Our previous mechanistic studies focused on the transcription factor (TF) NR2F1 and a handful of dormancy genes, revealing that the AZA+atRA reprogramming activated key dormancy pathways. However, we did not uncover the extent and functional characteristics of the programs activated by the reprogramming protocol, the NR2F1-independent programs that contributed to reprogramming into dormancy, and whether this strategy could indeed suppress metastatic progression *in vivo*. Mechanistic understanding of the AZA+atRA reprogramming is vital because this strategy, derived from our past work^14^, repurposed two FDA-approved drugs (e.g., AZA and atRA) to treat prostate cancer patients at risk of developing metastasis (clinicaltrials.gov identifier NCT03572387).

Here we provide a comprehensive understanding of the gene expression programs that epigenetic- and morphogen-driven reprogramming activates in HNSCC models. We reveal how the AZA+atRA reprogramming protocol, surprisingly, only taps into a sub-program of genes found in dormant cells; it identifies new programs not evident in spontaneous dormant cells and reveals how NR2F1-independent TGFβ-SMAD4-dependent mechanisms contribute to dormancy induced by AZA+atRA therapy. Importantly, we show that indeed AZA+atRA or AZA combined with AM80 (a clinically approved RARα agonist) used in a neo-adjuvant + adjuvant setting can significantly suppress aggressive HNSCC PDX metastasis to mouse lungs. This phenotype was associated with the induction of NR2F1 and the NR2F1-independent regulation of SMAD4 in DCCs. Our findings provide insight into the mechanisms behind the reprogramming of cancer cells using AZA+atRA, additional markers to pinpoint spontaneously occurring dormant cancer cells or DCCs in response to epigenetic therapies. Our data also support that even cancer cells with highly aberrant genomes can be reprogrammed into a dormant phenotype that may curtail metastatic progression.

## RESULTS

### Transcriptional programs controlled by AZA+atRA upon epigenetic reprogramming of HNSCC cells

We had previously shown that treatment of T-HEp3 HNSCC PDX cells and other HNSCC, prostate, and breast cancer cells with AZA+atRA *in vitro* could reprogram these cancer cells into quiescence when implanted *in vivo* and this phenotype correlated with the induction of dormancy genes such as NR2F1, SOX9, p21 and RARβ and a dormant phenotype sustainable for 2-4 weeks^14^. However, this analysis only provided a focused analysis of NR2F1 computationally predicted target genes^14^. Therefore, we sought to understand further the global gene expression programs and pathways regulated by AZA+atRA mediated epigenetic reprogramming, as a better understanding of these mechanisms may be helpful upon future use of the AZA+atRA or other epigenetic therapies.

To this end, we explored how the AZA+atRA reprogramming impacted the transcriptional landscape of T-HEp3 HNSCC malignant cells, compared to the programs activated in dormant D-HEp3 cells^17^. We treated T-HEp3 primary culture cells from growing PDX tumors sequentially with AZA (5 nM) and atRA (1 uM) and performed RNA sequencing (RNA-seq) **(Figure 1a)**. Principle component analysis (PCA) of the RNA sequencing data across samples showed that replicate samples for each condition are highly similar **(Figure S1a)**. Differential gene expression analysis (log2fc < or > 0, padj < 0.05) showed that 235 genes were upregulated, and 67 genes downregulated by AZA+atRA reprogramming compared to DMSO control **(Figure 1b, Table S1)**. Several well-known targets of retinoic acid signaling (STRA6, RARβ, HOXA5, and TIMP1) were upregulated^18, 19, 20^, and further validation by qPCR (**Figure S1b**) confirmed that the AZA+atRA mediated reprogramming in T-HEp3 cells was effective. Gene set enrichment analysis (GSEA) shows that the significantly enriched hallmark gene sets upregulated by AZA+atRA reprogramming are KRAS signaling, estrogen response early, mTORC1 signaling, and inflammatory response **(Figure S1c)**. In addition to these pathways, TGFβ receptor signaling is one of the significantly enriched pathways in the Wikipathways database **(Figure S1d)**. Many studies, including our previous reports, have shown that TGFβ/SMAD pathway plays an important role in normal stem cell and cancer cell dormancy, where a higher ratio of TGFβ2/TGFβ1 promotes dormancy via specific TGFβ receptors^21, 22, 23, 24^. Further, we performed ChIP Enrichment Analysis^25^ using the differentially expressed genes upon AZA+atRA reprogramming, and found a significant enrichment of SMAD2 and SMAD3, accounting for 21.7% (51/235) of differentially expressed up-regulated genes **(Figure 1c)**. We conclude that AZA+atRA may induce a SMAD-dependent program of growth arrest that may be associated with TGFβ2 induction^14^.

**Figure 1.**
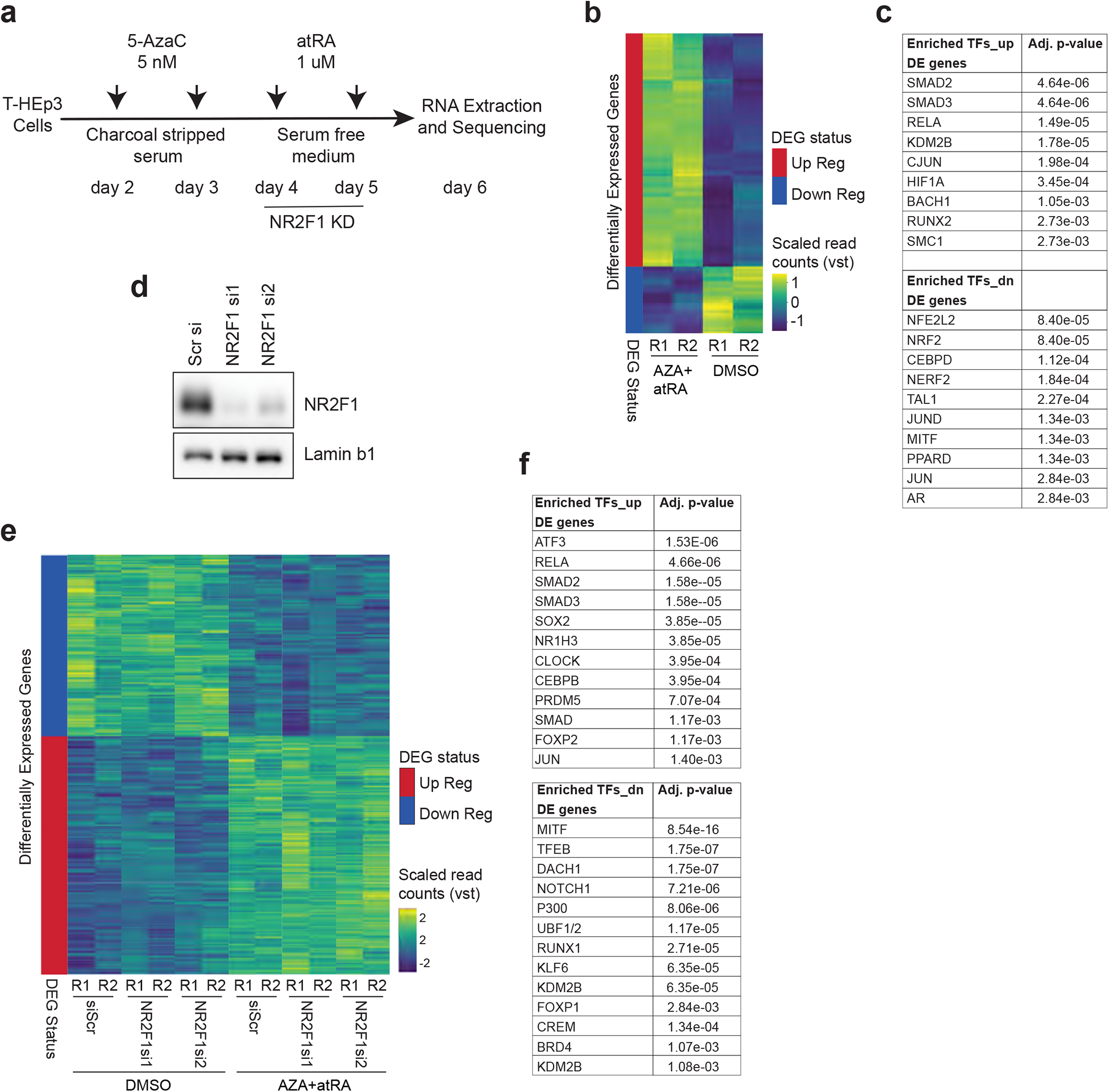
Transcriptional changes induced by AZA+atRA reprogramming. **a**. AZA+atRA reprogramming protocol. T-HEp3 cells seeded at low density were treated with AZA (5 nM) in DMEM medium containing charcoal stripped FBS for two days at the interval of 24 hrs. After 48 hours, AZA containing culture medium was replaced with serum-free DMEM medium, and cells were treated with atRA (1 uM) and NR2F1 knockdown was carried out simultaneously for 48 hours. Cells were collected and processed for RNA extraction, followed by RNA sequencing. **b**. Heatmap shows differentially expressed genes (DEGs) in AZA+atRA reprogrammed T-HEp3 cell line compared to DMSO control identified by RNA-seq analysis (2 biological replicates per condition). R: replicate. **c**. ChIP enrichment analysis (ChEA) of DEGs in Figure 1b using Enrichr to compute over-representation of transcription factor (TF) targets. Only significantly enriched TFs are shown. **d**. Western blot shows NR2F1 knockdown efficiency of siRNAs in T-HEp3 cells. **e**. Heatmap shows the comparison of DMSO control and AZA+atRA reprogramming irrespective of NR2F1 status (scr or NR2F1 knockdown) to identify DEGs independent of NR2F1. **f**. ChEA of DEGs in AZA+atRA reprogramming independent of NR2F1 using Enrichr to compute over-representation of transcription factor targets. Only significantly enriched TFs are shown.

### Epigenetic reprogramming with AZA+atRA induces NR2F1 dependent and independent transcriptional programs

We next sought to identify NR2F1 dependent and independent gene expression programs caused by AZA+atRA, we performed RNA sequencing in T-HEp3 cells depleted of NR2F1 by siRNAs and reprogrammed with AZA+atRA **(Figure 1a, d)**. Knockdown of NR2F1 by siRNAs in D-HEp3 and subsequent *in vivo* CAM assay led to a rapid switch from dormancy to the proliferative stage also confirmed the efficacy of siRNAs **(Figure S2a)**. Differential gene expression (DGE) analysis showed a small set of genes, 81 genes upregulated and 18 genes downregulated, dependent on NR2F1 in AZA+atRA reprogramming **(Figure S2b, Table S2)**. The small number of NR2F1-dependent (NR2F1-dep) DEGs revealed two significantly enriched pathways by AZA+atRA reprogramming in culture (hypoxia and IL6 JAK STAT3), hypoxia having been previously linked to NR2F1 induction^26^. Next, we set to identify genes responsive to AZA+atRA treatment but independent of NR2F1 **(Figure 1e)**. As expected, we found a higher number (943 downregulated and 1240 upregulated) of DEGs in AZA+atRA reprogramming, which were independent of NR2F1 expression **(Figure 1e, Table S3)**, arguing that while NR2F1 is important for the reprogramming, a larger program(s) is activated independently by this therapeutic approach. GSEA showed that the significantly enriched hallmark gene sets upregulated by AZA+atRA reprogramming in an NR2F1-independent (NR2F1-ind) manner were cholesterol homeostasis, mTORC1 signaling, epithelial-mesenchymal transition (EMT), and KRAS signaling while downregulated gene sets were enriched for MYC target and hypoxia **(Figure S2e)**. ChEA analysis of the DEGs in AZA+atRA reprogramming that are NR2F1-ind showed again enrichment of SMAD2, SMAD3 and SOX2 TFs **(Figure 1f)**. These data supported that, as hinted by the induction of TGFβ2 by AZA+atRA reported earlier^14^, activation of TGFβ-SMAD signaling may be important for AZA+atRA-induced dormancy.

### AZA+atRA induces a dormancy-like gene expression program that mainly overlaps with NR2F1-independent targets

To compare the similarity in gene expression signatures between the dormancy induced by AZA+atRA in T-HEp3 and dormancy programs in D-HEp3 cells, we performed RNA-seq comparing T-HEp3 and D-HEp3 cells. *In vivo* proliferative T-HEp3 primary culture cells at passage 0 (P0) and *in vivo* dormancy D-HEp3 passage >130 as reported previously^14^, were used for the RNA-seq experiment. Analysis of the RNA-seq data revealed that 3597 and 3451 genes were differentially upregulated in T-HEp3 and D-HEp3, respectively **(Figure 2a)**. Confirming the phenotypes, some of the critical genes previously found upregulated in D-HEp3 cells and described by microarrays data^14^ were also found in the RNA-seq, and this included TGFβ2, NR2F1, SOX9, LIF, and KLF4, while in T-HEp3 cells, MMPs (1, 2, 3, 8, 9, 13, 17, and 19), VIM, VEGFA, IL1B, IL24, IL8, and JUN were upregulated **(Figure 2a, Table S4)**. GSEA of DEGs showed that genes upregulated in D-HEp3 were enriched for hallmark pathway E2F targets, mTORC1 signaling, Myc targets, G2M checkpoint, oxidative phosphorylation, glycolysis and, fatty acid metabolism, consistent with their growth suppression but adaptation to survive *in vivo* **(Figure S3a)**. Upregulated genes in T-HEp3 cells were enriched for hallmark pathways such as EMT, TNFA signaling, KRAS signaling, IL2 STAT5 signaling, inflammatory response, angiogenesis, and myogenesis **(Figure S3a)**. ChEA of DEGs in D-HEp3 and T-HEp3 did not show significant enrichment of SMAD2/3 target genes (adjusted p-value were higher than 0.05) **(Figure S3b)**. Since AZA+atRA reprogrammed T-HEp3 cells displayed growth suppression^14^, we compared the DEGs of AZA+atRA reprogramming with the DEGs of the D-HEp3 cell line. We observed 27 (11.48%) upregulated and 37 (55.22%) downregulated genes (both NR2F1-dep and NR2F1-ind) of AZA+atRA reprogrammed T-HEp3 cells overlapped with upregulated and downregulated genes, respectively in spontaneously dormant D-HEp3 cells **(Figure S3c)**. Some of the genes that showed a positive correlation between dormant D-HEp3 cells and AZA+atRA reprogrammed T-HEp3 were STRA6, LIF, KLF4, HOXD3, COL4A1, IL24, and FOXQ1 **(Table S5)**. Next, we checked the NR2F1-ind DEGs of AZA+atRA reprogramming, and differentially expressed in D-HEp3. A total of 450 (20.61%) DEGs of AZA+atRA reprogramming overlap with DEGs of D-HEp3, which accounts for 181 (14.59%) upregulated and 269 (28.52%) downregulated genes **(Figure 2b, Table S6)**. However, the data revealed that ∼80% of the genes induced and repressed by the AZA+atRA reprogramming of T-HEp3 cells did not match those regulated in D-HEp3 cells, arguing for a unique program activated by this drug and morphogen combination. Thus, the AZA+atRA reprogramming (both NR2F1-dep and NR2F1-ind) strategy only taps into a subprogram found in spontaneously dormant D-HEp3 cells, and that other AZA+atRA specific gene programs may also be able to cooperate with the changes observed in 20% of genes in D-HEp3 cells to induce a strong and long-lived growth suppression.

**Figure 2.**
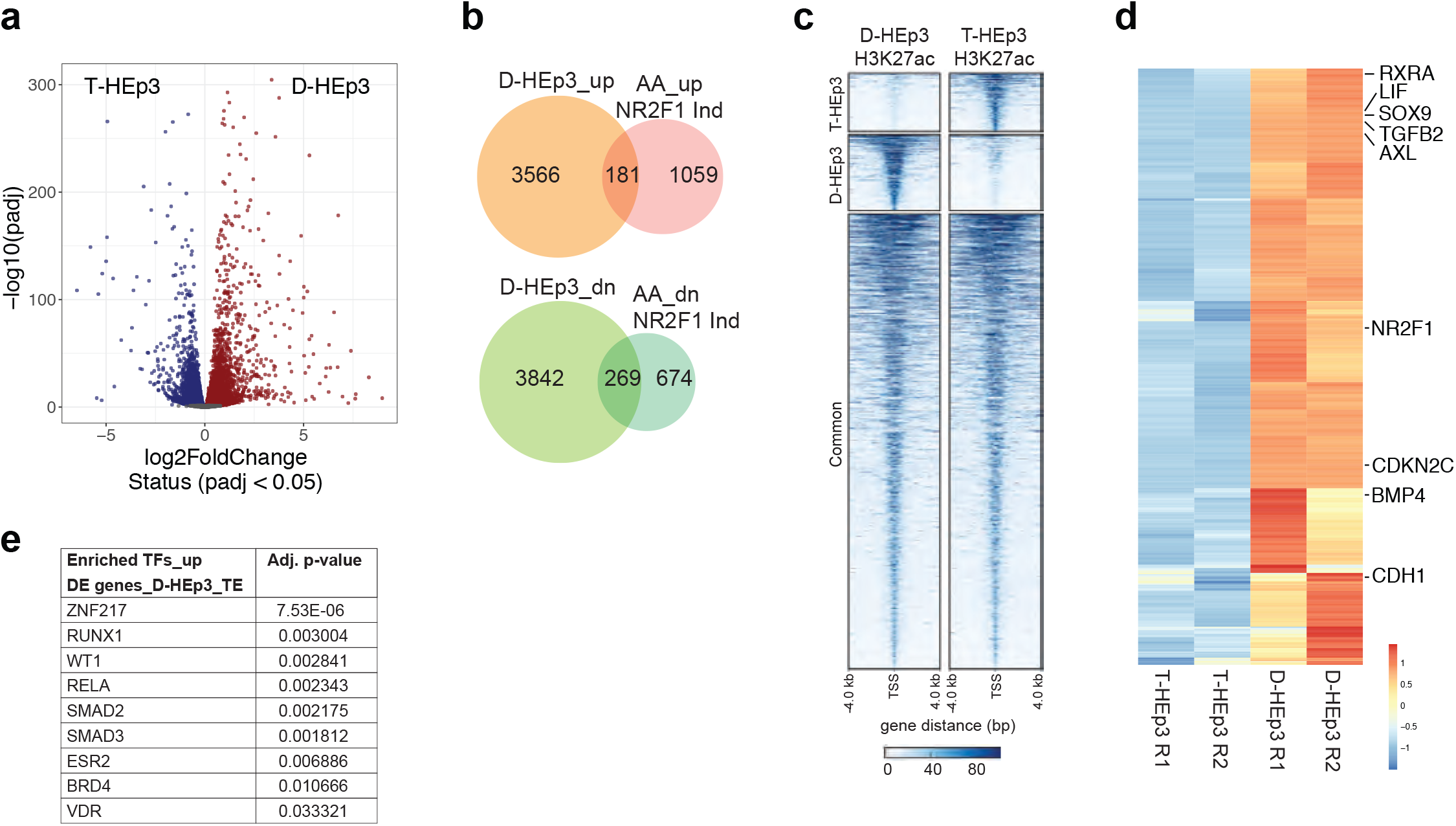
RNA-seq analysis of D-HEp3 and T-HEp3 transcriptome. **a**. Volcano plot showing the DEGs (upregulated) in D-HEp3 (red) and T-HEp3 (blue). Adjusted p-value (≤ 0.05) was used as a cut-off to identify DEGs. **b**. Venn diagram shows the comparison of upregulated and downregulated genes in D-HEp3 with up and downregulated NR2F1-independent genes upon AZA+atRA reprogramming. **c**. ChIP-seq analysis of H3K27ac marks was performed in D-HEp3 and T-HEp3 cells. Enriched peaks (unique to either D-HEp3 or T-HEp3 and common to both D-HEp3 and T-HEp3) are shown as density plot. **d**. DEGs in D-HEp3 was overlaid with H3K27ac enriched marks in D-HEp3, and the heatmap shows upregulated genes in D-HEp3 regulated by enhancers. **e**. ChEA of enhancer regulated genes in D-HEp3 using Enrichr to compute over-representation of transcription factor targets.

### AZA+atRA epigenetic reprogramming dependent program of dormancy involves the SMAD pathway

To further explore whether TGFβ-SMAD signaling could be a pathway activated epigenetically, we focused on enhancer activity^27, 28^. The gain and loss of the activity of these enhancers plays an important role in metastatic progression. The activity of enhancers is defined by specific chromatin signatures like histone modifications (H3K4me1 and H3K27ac) and DNase-hypersensitivity. To this end, we conducted chromatin immunoprecipitation coupled to high-throughput sequencing (ChIP-seq) for H3K27ac as a marker of active enhancers in T- and D-HEp3 cells^29, 30^. We identified 2461 and 3177 differential enhancers in T- and D-HEp3 cells, respectively **(Figure 2c)**. Next, differentially enriched distal/intragenic H3K27ac regions were associated to the positively correlated promoters of DEGs in D-HEp3 vs. T-HEp3 cells within a genomic region of ± 500 kb and identified 589 and 703 genes regulated by enhancers, respectively **(Figure 2d, S3d, Table S7 and S8)**. This analysis revealed that dormancy-associated genes upregulated in D-HEp3, such as TGFβ2, NR2F1, SOX9, and LIF, were associated with enhancers. We next performed ChEA in genes regulated by enhancers in D-HEp3 and observed a significant enrichment of genes regulated by SMAD2 and SMAD3 **(Figure 2e)**. Our analysis showed enrichment of SMAD2 and SMAD3 target genes in AZA+atRA reprogramming and enhancer-regulated genes in D-HEp3 **(Figure S3e)**, suggesting a prominent role of the SMAD pathway in D-HEp3 cells’ dormancy. This result is in line with our previous study, where we showed the role of TGFβ2 and TGFβRIII in the regulation of DCCs fate in different organs ^24^.

### Restoration of TGFβ and SMAD4 signaling independently of NR2F1 is a hallmark of AZA+atRA reprogramming into dormancy

The results revealing a SMAD2/3 signaling signature suggested that the AZA+atRA reprogramming strategy could be restoring the canonical growth-suppressive function of the TGFβ pathway active during the early stages of cancer progression that then shows loss of SMAD4 function^31^.

To explore this possibility, we first tested the expression of SMAD (2, 3, and 4) proteins in AZA+atRA reprogrammed T-HEp3 cells. We observed that all three SMADs are, as expected, low in malignant T-HEp3 cells, and AZA+atRA treatment caused a significant increase in the total expression level of SMAD2, SMAD3, and SMAD4 **(Figure 3a)**. We had published that TGFβ2 induced DCC dormancy in the bone marrow *via* an activated P-p38 as well as SMAD2 and SMAD1/5 pathway, which induced expression of p27 leading to quiescence^23, 24^. Upon AZA+atRA reprogramming, we observed an increase in the mRNA levels of TGFβ2^14^, along with TGFβ1 and TGFβ3 (**Figure S4a)**. However, we did not observe any considerable change in the expression or phosphorylation of the SMAD1/5 pathway as reported for TGFβ2^24^ **(Figure 3a)**, arguing that the AZA+atRA reprogramming may induce a canonical pathway activation. We also documented an increase in the expression of TGFβRI and TGFβRIII **(Figure S4b)**, both previously implicated in DCC dormancy in the BM and lung^23, 24^. Upon AZA+atRA reprogramming, we also detected a strong increase in the phosphor-p38 and p27, while P-Erk1/2 or total Erk1/2 remained constant, leading to a low ERK/p38 activity ratio^14, 32^ **(Figure 3a)**. In contrast to what we detected in D-HEp3 cells^24^, TGFβ1 treatment of AZA+atRA reprogrammed T-HEp3 cells, which already expresses high TGFβ1 levels, did not cause their re-awakening *in vivo* (data not shown), suggesting a change in TGFβ1 signaling pathway.

**Figure 3.**
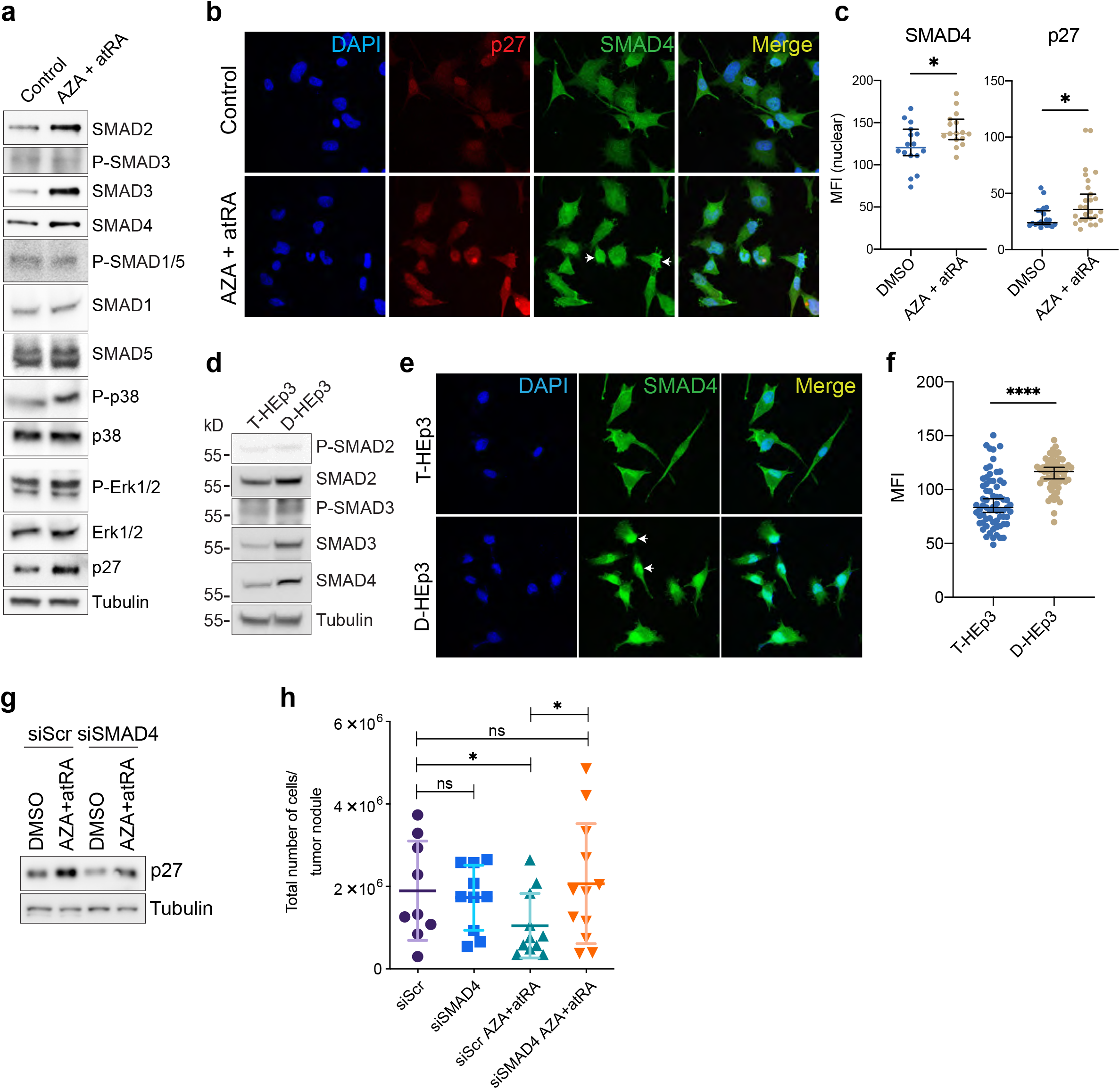
AZA+atRA reprogramming induces SMAD2, SMAD3, and SMAD4 expression. **a**. Western blots show upregulation of SMAD2/3/4, dormancy associated marker genes like p27 and P-p38 upon AZA+atRA reprogramming in T-HEp3 cells. **b**. Immunofluorescence (IF) staining show the higher nuclear localization of p27 and SMAD4 (marked with solid white arrowheads) in AZA+atRA reprogrammed T-HEp3 cells. **c**. Quantification of nuclear SMAD4 and p27 in control and AZA+atRA reprogrammed T-HEp3 cells (unpaired t-test, p ≤ 0.05). **d**. Western blots show higher expression of SMAD2/3/4 in D-HEp3 cells compared to T-HEp3 cells. **e**. IF staining shows higher SMAD4 nuclear localization (marked with solid white arrowheads) in D-HEp3 cells. **f**. Quantification of nuclear SMAD4 in D-Hep3 and T-HEp3 cells (unpaired t-test, p ≤ 0.05). **g**. SMAD4 knockdown by siRNA and AZA+atRA reprogramming in T-HEp3 cells reveals p27 dependency on SMAD4. **h**. T-HEp3 (1.5 × 10^5^) cells depleted of SMAD4 and AZA+atRA reprogrammed were inoculated in chicken CAM and incubated for four days. After incubation, tumors were excised, and the total number of cells counted (experiment was repeated three time with a minimum of three eggs per experiment per condition, Mann-Whitney test, p ≤ 0.05).

SMAD proteins frequently shuttle between the cytoplasm and the nucleus and control transcription of target genes, but the basal level of SMAD proteins in the cytoplasm and the nucleus remains constant^33, 34^. Therefore, since we observed an increased expression of SMAD2/3/4, we anticipated their higher nuclear localization for transcriptional activation of target genes. To this end, we reprogrammed T-HEp3 cells with AZA+atRA and performed SMAD4 and p27 immunofluorescence (IF) and immunoblot assay; we observed an increased nuclear localization for SMAD4 and p27 **(Figure 3b-c, S4c)**. Similarly, reprogramming of T-HEp3 cells with AZA+AM80 (RARα agonist) (50 nM) also led to an increase in SMAD4 and p27 levels **(Figure S4c-e)**. These data show that the AZA+atRA reprogramming orchestrates a restoration of TGFβ/SMAD signaling, notably with upregulation of SMAD4, in malignant HNSCC cells.

To determine whether AZA+atRA was activating a TGFβ pathway as observed in dormant D-HEp3 cells, we detected the expression level of SMAD proteins in T- and D-HEp3 cells. We observed significantly higher expression of SMAD2, SMAD3, and SMAD4 in D-HEp3 cells **(Figure 3d)**. In addition to higher expression, there were more nuclear-localized SMAD4 in D-HEp3 cells **(Figure 3e-f)**. Knockdown of SMAD4 was sufficient to reverse the AZA+atRA induction of p27 in T-HEp3 cells and it partially reversed AZA+atRA-induced growth suppression of T-HEp3 cells on the CAM *in vivo* assay **(Figure 3g-h, S4f)**. These data support that AZA+atRA reprogramming induces, independently of NR2F1, a dormancy-like phenotype in proliferative cancer cells in part by activating a TGFβ-p38-SMAD4 pathway. The data also support that the AZA+atRA reprogramming protocol is robust in inducing growth suppression because it taps into differentiation (retinoic acid)- and dormancy-inducing (TGFβ2) pathways.

### AZA+atRA or AZA+AM80 induces dormancy of NR2F1+/SMAD4+ DCCs suppressing metastatic outgrowth in the lungs

Our data support that the epigenetic and transcriptional changes in reprogrammed cancer cells via the AZA+atRA treatments, which has only been tested for short periods in primary site tumor growth^14^, could induce a long-lived and self-sustained program of canonical TGFβ-SMAD2/3/4 signaling that could limit the metastatic expansion of pre-existing DCCs.

To this end, we tested the effect of AZA treatment followed by atRA or AM80 (a RARα agonist with a longer half-life than atRA) on metastatic progression. Immunocompromised nude mice (BALB/c nu/nu) were injected s.c. with 0.5 × 10^6^ T-HEp3-GFP cells obtained directly from the PDX maintained tumors^35^. After primary tumors reached a size of 300 mm^3^, mice were treated with vehicle or with one cycle of neoadjuvant (AZA+atRA or AZA+AM80); AM80 is an FDA-approved RARα-specific agonist used in hematological malignancies treatment^36^. We did this neoadjuvant step because T-HEp3 cells can disseminate very early after implantation already seeding lungs when tumors are smaller than 300 mm^3^ ^24^. We used 1 mg/kg/day of AZA and atRA and 0.3 mg/kg/day of AM80 doses following the previous optimization of the dosing ^37^. After the neoadjuvant cycle, tumors (600 mm^3^) were removed by surgery, and after 48 hours, mice were treated for four adjuvant cycles with the vehicle, AZA+atRA or AZA+AM80. At the end of the fourth cycle, mice were euthanized, and the number of T-HEp3-GFP positive tumor cells were scored in one lung lobule enzymatically digested with collagenase-I **(Figure 4a)**. These fresh collagenase-I suspensions already scored for GFP+ events were further stained in suspension using anti-vimentin human-specific antibodies as reported previously^14^ to confirm the GFP counts. This second quantification was included because we previously reported that dormant DCCs could lower GFP expression, preventing the proper detection of quiescent cells^24^. The other lung lobule was processed for histological and immunofluorescence analysis for further confirmatory analysis of DCC phenotypes. The neoadjuvant treatment had no significant effect on primary tumor growth **(data not shown)**. However, compared to the control vehicle, mice treated with AZA+AM80 showed a significant decrease in the number of T-HEp3-GFP cells in the lungs **(Figure 4b)**. AZA+atRA also showed a decrease in the DCCs but not as effective as AM80. Quantification of the same samples but for the number of vimentin-positive DCCs, confirmed the GFP count differences between groups but revealed that both AZA+atRA or AZA+AM80 treated mice showed a significant decrease in tumor cells burden in the lungs **(Figure 4c-d)**. The suppression of metastatic burden correlated with dormancy induction as protein expression of NR2F1 was significantly upregulated in the DCCs and micro-metastasis in the lungs of mice treated with AZA+atRA or AZA+AM80 **(Figure 4e)**. We conclude that a 5-week neoadjuvant+adjuvant treatment of mice with already existing metastatic spread can significantly suppress metastatic progression, which correlated with the induction of the dormancy regulator NR2F1, confirming the activation of the NR2F1-dependent program.

**Figure 4.**
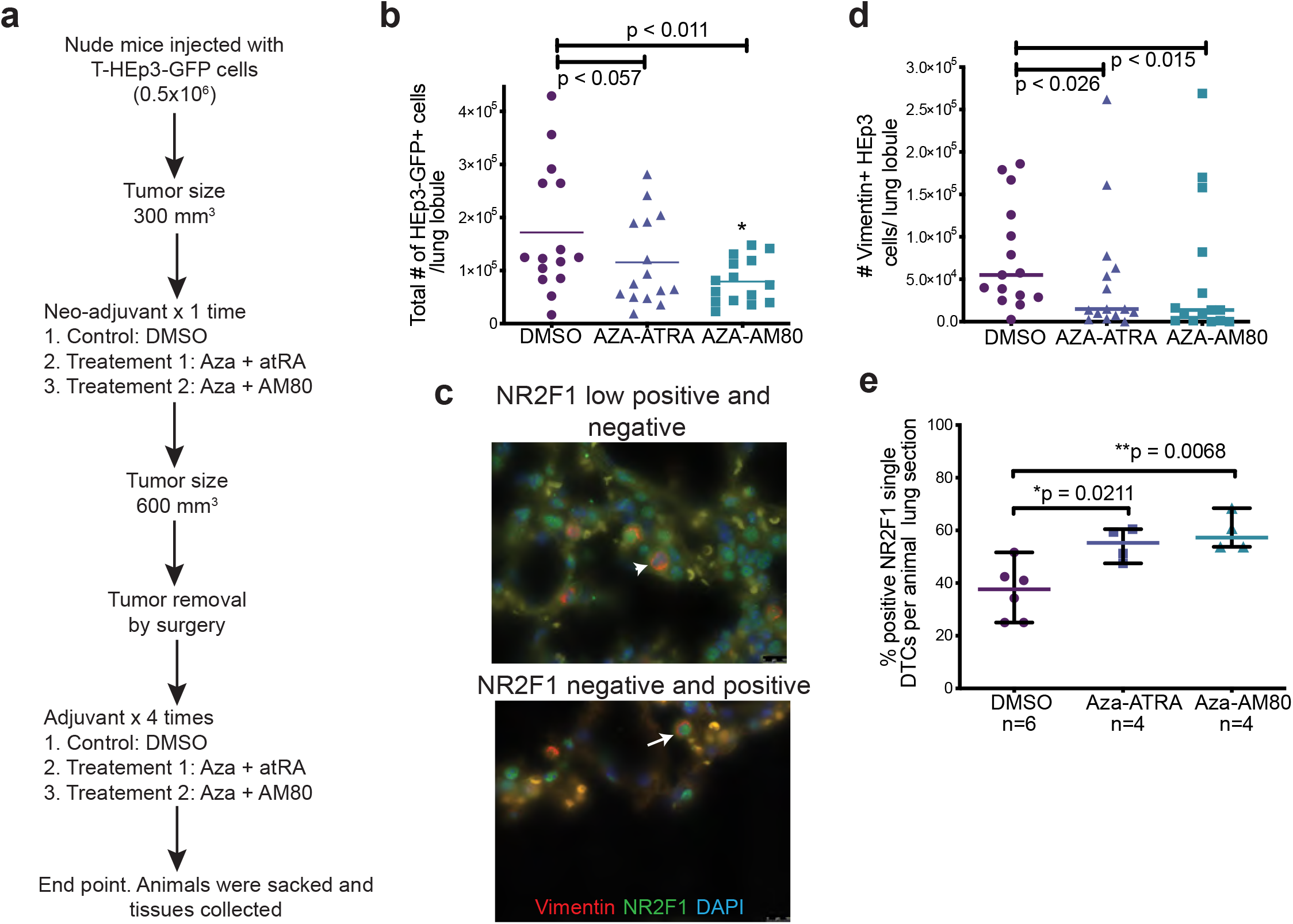
AZA+atRA and AZA+AM80 suppress the metastatic load in an experimental mouse model. **a**. Schematic of *in vivo* mouse experiment. 0.5 × 10^6^ T-HEp3-GFP primary tumor cells (passage 0 from CAM) were injected into nude mice. Mice were treated in neo-adjuvant setting with AZA+atRA or AZA+AM80 once when tumor size reached an average size of 300 mm^3^. At 600 mm^3^ of tumor size, surgery was performed to remove the primary tumor from the animals and treated four cycles (28 days) of adjuvant therapy (AZA+atRA or AZA+AM80). Animals were euthanized and organs processed for analysis. **b**. Mouse lungs digested enzymatically and counted for GFP+ T-HEp3 single cells. **c**. T-HEp3-GFP cells in the digested lung suspensions were stained with vimentin and NR2F1. Representative images show positive (white arrow) and negative signal (white arrowhead) for NR2F1. **d**. Quantification of T-HEp3-GFP cells in lung sections using vimentin IF staining. **e**. Quantification of NR2F1+ and vimentin+ T-HEp3-GFP cells in lung sections.

Next, we tested whether the SMAD4 program could inform on HNSCC patient outcome. To this end we tested whether expression of SMAD2/3 and SMAD4 target genes in primary tumors associated with differences in disease free survival^38^. We found that patients with enrichment for the SMAD2/3 or SMAD4 signatures relapsed at a slower rate than patients low for this signature, albeit with limited statistical significance **(Figure S5)**. This suggests that DCCs derived from tumors with a high SMAD signature may take longer to activate from dormancy. Because access to patient’s target organs to find DCCs was not available we tested this possibility in mice in control or under AZA+atRA treatments. When we tested whether the SMAD4 program could be detected in the mouse lungs treated with vehicle control, AZA+atRA, or AZA+AM80 we found that SMAD4 was upregulated spontaneously in solitary DCCs compared to micro- and macrometastatic lesions that had very low or no expression of SMAD4 **(Figure 5a-d)**. This argues that spontaneously dormant DCCs turn on the SMAD4 program. Macrometastasis were not detected in the lungs of mice treated with AZA+atRA or AZA+AM80 groups supporting a powerful metastasis suppressive activity. In the AZA+atRA group, while no micro- or macro-metastasis were found, even the solitary residual DCCs expressed higher levels of SMAD4. In the AM80 group, solitary DCCs displayed similar levels of SMAD4 as the solitary DCCs in the control group, but the few micro-metastases in the AM80 group displayed significant upregulation of SMAD4 compared to metastasis in the control group **(Figure 5e)**.

**Figure 5.**
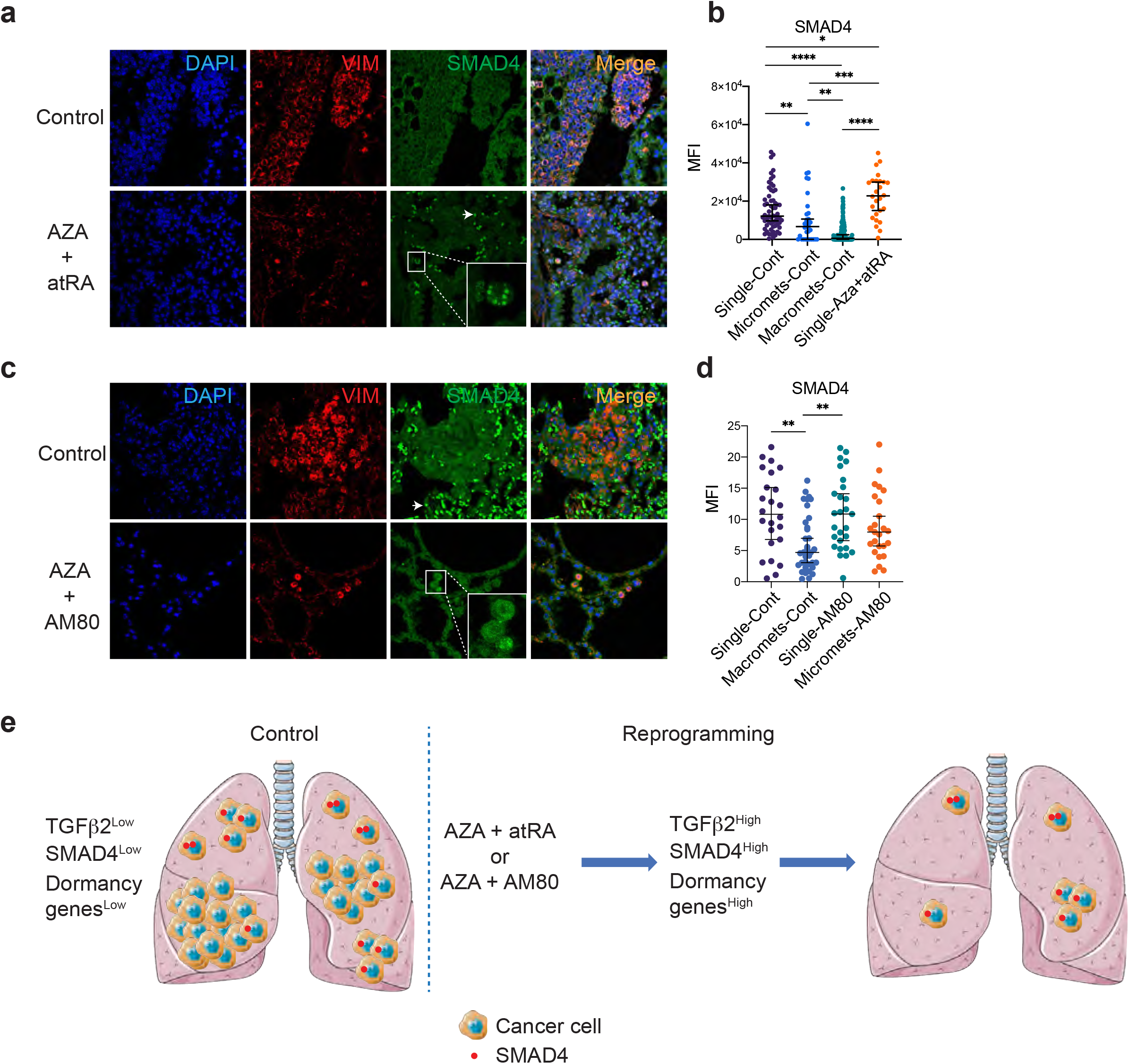
SMAD4 keeps DCCs in a dormant single-cell state. **a**. IF staining images showing SMAD4 expression in disseminated T-HEp3-GFP cells in control and AZA+atRA treated mouse lungs. **b**. Quantifications of mean fluorescent intensity (MFI) of SMAD4 signals in T-HEp3-GFP cells in control and AZA+atRA treated mouse lungs (Mann-Whitney test, p ≤ 0.05). **c**. IF staining images showing SMAD4 expression in disseminated T-HEp3-GFP cells in control and AZA+AM80 treated mouse lungs. **d**. Quantifications of MFI of SMAD4 signals in T-HEp3-GFP cells in control and AZA+AM80 treated mouse lungs (Mann-Whitney test, p ≤ 0.05). **e**. Working model: lungs of control animals harbor macrometastasis, micrometastasis, and single DCCs, while treatment with AZA+atRA or AZA+AM80 inhibits metastatic growth leading to micrometastasis and single DCCs. Single DCCs have higher SMAD4 expression which keeps cancer cells in a dormant single-cell state.

## DISCUSSION

Our results produced a series of findings that warrant further discussion. Our first goal was to better define the gene programs activated by the AZA+atRA reprogramming protocol that we had discovered can turn on dormancy programs when following a computationally predicted signature of NR2F1 targets. Our second goal was to test whether indeed such reprogramming therapy could suppress metastasis *via* induction of a dormancy-like program. Both goals were achieved, and the data revealed informative differences between therapy-induced and spontaneously activated dormancy pathways.

Analysis of the AZA+atRA-induced gene programs (both NR2F1-dep and NR2F1-ind) and those found in spontaneously dormant HEp3 cells showed that the reprogramming therapy only regulates a subset and rather small subprogram found in dormant D-HEp3 cells. However, this program seems to be strong enough to suppress metastasis development, at least for the time the therapy is administered. Our data also revealed that an even smaller sub-program of gene regulation is dependent on NR2F1 upon AZA+atRA reprogramming of malignant HEp3 cells, which again is important for the growth suppressive effect of the AZA+atRA treatment. We found that multiple subprograms are activated when we focused on the larger NR2F1-independent mechanism, but intriguingly a TGFβ-SMAD program was evident. This program was also found in D-HEp3 cells and has been revealed in other models as well^24^. Components of the TGFβ-SMAD program seems to be epigenetically controlled as our analysis revealed that genes controlled by specific enhancers in D-HEp3 cells also show enrichment for SMAD2/3 regulated genes. This program we have shown is associated with TGFβ2 but not TGFβ1 signaling, the latter being crucial for the reactivation of these cancer cells^24^. However, the AZA+atRA-induced reprogramming seemed to reverse the pro-growth function of TGFβ1 and restore canonical signaling for this pathway. In fact, TGFβ1 is not a reactivation signal in AZA+atRA-induced dormant T-HEp3 cells. AZA+atRA resulted in the upregulation of all TGFβ ligands and TGFβRI and TGFβRIII and strong upregulation of SMADs2, 3, and 4. The latter is commonly silenced or mutated in HNSCC and other cancers^39, 40^, and the strong upregulation found in the *in vitro* reprogramming, but also in the solitary DCCs in lungs was remarkable. This result suggested that AZA+atRA may restore to some extent the tumor-suppressive function of the TGFβ pathway even in HNSCC PDX tumors with highly aberrant genomes. SMAD4 induction seems to be necessary for induction of p27 and also for growth arrest upon AZA+atRA treatment; however, it is not the only pathway that may be contributing as many other pathways and transcription factors seem to be contributing, albeit partially, to the growth arrest program.

The multiplicity of programs, NR2F1-dependent and independent, activated by the AZA+atRA treatment may explain the long-lived and durable effect of the reprogramming observed in primary sites^14^, bone-marrow of breast cancer patients^41^ and metastasis in this study. Dosing of AZA+atRA using a neo-adjuvant+adjuvant scheme was effective in suppressing metastasis and DCCs upregulated both NR2F1 and SMAD4, supporting that the programs revealed in the RNA-seq are detected in the growth arrested DCCs. Notably, the use of these two proteins as biomarkers may help pinpoint spontaneously dormant DCCs or those induced by epigenetic therapies. Similarly, perhaps the use of an NR2F1 and/or SMAD4 signature in primary tumors may already identify patients with different risk levels for relapse. Interestingly, the clinically approved agonist for RARα was also effective in inducing metastasis suppression after the AZA treatment. This may be important for future clinical applications as the AZA+atRA reprogramming protocol is being used in a prostate cancer clinical trial, and more potent agonists of RARα may be useful to treat these patients. In addition, we have also found that a novel NR2F1 agonist can suppress metastasis^42^. Interestingly, a neural crest differentiation program was activated by the NR2F1 agonist^42, 43^ which we did not find with the AZA+atRA protocol. The NR2F1 agonist did not activate the TGFβ pathway^42^, in agreement with our findings in this study that this is an NR2F1 independent mechanism.

While we did not test the activation or silencing of enhancers with AZA+atRA, the presence of active SMAD2/3-linked enhancers in D-HEp3 cells suggests that chromatin remodeling and epigenetic reprogramming even of complex genomes may be sufficient to suppress metastatic potential as observed in the transition of T-HEp3 cells to D-HEp3 cells or AZA+atRA-reprogrammed cancer cells. Further studies will explore how enhancers and super-enhancers revealed in our analysis contribute to the stability and durability of the dormancy program and whether they may serve as biomarkers of a durable dormancy phenotype.

Our new data supports that therapeutically induced <u>d</u>ormant <u>c</u>ancer <u>c</u>ells or iDCCs may represent an alternative for managing residual latent cancer or progressive disease that may not be manageable with standard care approaches. The latter may be the case in an ongoing clinical trial in prostate cancer (clinicaltrials.gov identifier NCT03572387) that employs AZA+atRA iDCCs to prevent the progression of biochemically relapsing disease. Our studies paired with the clinical trial results may provide important information on how to optimize iDCC therapies in the near future.

## ACKNOWLEDGEMENTS

We thank the Aguirre-Ghiso, Bernstein and Sosa labs for helpful discussions. This work was supported by grants from National Institutes of Health (NIH)/National Cancer Institute (NCI) (CA109182, CA216248, CA218024, CA196521 to JAAG and RO1CA154683, CA218024 to EB) and The Jimmy V Foundation to JAAG. JAAG is Samuel Waxman Cancer Research Foundation Investigator. MSS was supported by Melanoma Research Foundation Career Development Award, NIH/NCI K22CA201054 and Melanoma Research Alliance team Science Award. RNA-seq and ChIP-seq analysis was supported in part through the Oncological Sciences Sequencing Core supported by Tisch Cancer Institute of the Icahn School of Medicine at Mount Sinai (ISMMS) Cancer Center Support Grant P30 (CA196521), Scientific Computing supported by the Office of Research Infrastructure of the NIH under award number S10OD026880 to ISMMS, and the Mount Sinai Genomics Technology Facility, and Deniz Demircioglu from the Bioinformatics for Next Generation Sequencing (BiNGS) shared resource facility within the Tisch Cancer Institute at the Icahn School of Medicine at Mount Sinai. The development of this shared resource is partially supported by the NCI P30 Cancer Center support grant.

## AUTHOR CONTRIBUTIONS

DKS and EF planned and conducted experiments, analyzed data, and wrote the manuscript; EF, JC, ARN and NK performed *in vivo* mouse experiments and DKS further processed *in vivo* material; DKS, SC, DH and DS performed RNA-seq and ChIP-seq experiments and analyzed data; JAAG, EB, DKS and MSS analyzed data, provided insight and wrote the manuscript. JAAG conceived the project with MSS and designed experiments along with MSS, DKS and EF. JAAG, MSS and EB secured funding.

## DECLARATION OF INTERESTS

JAAG is a scientific co-founder of, scientific advisory board member and equity owner in HiberCell and receives financial compensation as a consultant for HiberCell, a Mount Sinai spin-off company focused on therapeutics that prevent or delay cancer recurrence.

## METHODS

### Cell lines and culture

Tumorigenic (T-HEP3) cells were derived from a lymph node metastasis in head and neck squamous cell carcinoma (HNSCC) patient and maintained as patient-derived xenograft on chick chorioallantoic membrane (CAM) as described previously^17, 44^. Dormant (D-HEp3) cells were obtained by *in vitro* passaging of T-HEp3 cells for more than 40 generations. Both cell lines were cultured in Dulbecco’s Modified Eagle Medium (DMEM) with 10% heat-inactivated fetal bovine serum (FBS) and 100U penicillin/0.1 mg/ml streptomycin. Cells were incubated and grown at 5% CO2 and 37°C. Cells were routinely tested for mycoplasma (PCR Mycoplasma test kit PK-CA91-1096, PromoCell).

### Aza and atRA reprogramming

T-HEp3 cells (P1) growing in a dish were treated with 5 nM 5-Azacytidine (Sigma #A2385) in DMEM containing charcoal-stripped 10% FBS. After 24 hr, AZA was replenished and allowed to grow for another 24 hrs. Next, AZA-containing medium was removed, and cells were treated with 1 µM all-trans retinoic acid (Sigma #R2625) in serum-free medium and allowed to grow for another 48 hrs.

### RNA sequencing and analysis

Total RNA from T-HEp3 cells reprogrammed with AZA+atRA was extracted using RNeasy mini kit. Library preparation and RNA sequencing were performed at GENEWIZ, LLC. (South Plainfield, NJ, USA). In addition, library preparation and RNA sequencing for D-HEp3 and T-HEp3 cell lines were performed at the Genomics Core Facility at the Icahn School of Medicine at Mount Sinai. A detailed method of RNA sequencing and analysis was described previously^42^.

### ChIP-sequencing

H3K27ac native ChIP was performed in D-HEp3 and T-HEp3 cells following the method previously described^45^. Reads were aligned to the human reference genome hg19 using Bowtie v1.1.2 (Langmead et al., 2009) with parameters –l 65 –n 2 –best –k 1 –m 1, and reads quality was assessed using fastQC. Duplicate reads were removed with PICARD v2.2.4 (Broad Institute). Binary alignment maps (BAM) files were generated with samtools v1.9^46^ and were used in downstream analysis. MACS2 v2.1.0^47^ was used to called significant peaks. Peaks within ENCODE blacklisted regions were removed. Coverage tracks were generated from BAM files using deepTools 3.2.1^48^ bam coverage with parameters– normalize using RPKM–bin size 10. For genomic annotation promoters (−4 kb to +4 kb) relative to the TSS were defined according to the human hg19 genome version. Heatmaps of genomic regions were generated with deepTools 3.2.1. The command compute matrix was used to calculate scores at genomic regions and generate a matrix file to use with plot heatmap, to generate plots.

### ChIP-seq differential enrichment analysis

For D-HEp3 and T-HEp3 H3K27ac ChIP-seq, the BAM files were merged in a single BAM and significant peaks were called using MACS2 narrow Peak 2.1.0 to generate a bed file with common set of regions. The BAM file of all the common regions was used to call enhancers using the Ranking Ordering of Super Enhancers algorithm (ROSE) and the BAM files of the T-HEp3 and D-HEp3 were used to map their specific reads the ROSE called enhancers. Differential enhancers were called If the log2 fold change was greater than 1.5.

### RNA interference

Gene-specific siRNAs (listed in supplementary information **Table 1**) were transfected twice to HEp3 cells at a final concentration of 50 nM with a gap of 24 hours, using Lipofectamine RNAiMax reagent (Invitrogen #13778075).

### Western blot

Cells were lysed in RIPA buffer, and protein concentrations were calculated using Pierce BCA protein assay kit (Thermo Scientific #23227) and a standard BSA curve. Samples were boiled in 4X laemmli sample buffer (Bio-Rad #1610747) for 10 minutes at 95°C. To prepare nuclear protein extracts for immunoblotting, NE-PER nuclear and cytoplasmic extraction reagents (Thermo Fisher # 78833) was used. 10-12% SDS–PAGE gels were run in running buffer (25 mM Tris, 190 mM glycine, 0.1% SDS) and transferred to PVDF membranes in transfer buffer (25 mM Tris, 190 mM glycine, 20% methanol). Membranes were blocked in 5% milk or BSA in TBST (Tris-buffered saline with 0.1% Tween-20) buffer. Membranes were incubated with primary antibodies overnight at 4°C. Membranes were washed in TBST buffer and incubated with HRP-conjugated secondary antibodies at room temperature for 1 hour. Western blots were developed using Amersham ECL Western Blot Detection (GE #RPN 2106) and GE ImageQuant LAS 4010. The antibodies used are listed in supplementary information **Table 2**.

### Quantitative PCR

RNA was extracted using RNeasy mini kit (Qiagen #74104), and cDNA was synthesized using M-MuLV reverse transcriptase enzyme (New England Biolabs #M0253S) following the manufacturer’s instructions. Real-time quantitative PCR was performed using PowerUP SYBR Green Master Mix (Applied Biosystems #A25741). Primers used for amplification are listed in supplementary information **Table 3**.

### Xenograft mouse model study

Use of female nude mice and experimental procedures were approved by the Institutional Animal Care and Use Committee (IACUC) of Icahn School of Medicine at Mount Sinai. 0.5 × 10^6^ T-HEp3-GFP cells were injected subcutaneously in 8-week old female BALB/c nu/nu mice (Jackson Laboratories). Mice were inspected regularly for arising tumors and when tumors reached ∼300 mm^3^, mice were treated with vehicle or one cycle of neoadjuvant AZA+atRA or AZA+AM80. 1 mg/kg/day of AZA (Sigma #A2385) and atRA (Sigma #R2625) and 0.3 mg/kg/day of AM80 (Tocris #3507) doses were used for each treatment cycle (2 days of AZA + 3 days of atRA or AM80 + 2 days rest). After the neoadjuvant cycle, mice were injected with anesthetics ketamine (80–120 mg) and xylazine (5 mg), and tumors (∼600 mm^3^) were removed by surgery. Forty-eight hours post-surgery, mice were treated for four adjuvant cycles with the vehicle, AZA+atRA or AZA+AM80. At the end of the fourth cycle, mice were sacrificed by euthanization, and lungs were collected.

### Immunofluorescence

Paraffin-embedded tissue sections were incubated in xylene (10 minutes twice) followed by graded ethanol rehydration (3 minutes each). Antigen retrieval for mouse lung tissues was performed in 10 mM citrate buffer (pH 6) for 40 minutes using a steamer. Tissues were permeabilized in 0.3% TritonX-100 + PBS for 10 minutes. Cultured cells were fixed in 4% formaldehyde on ice for 10 minutes. Tissue sections and cultured cells were blocked with 3% bovine serum albumin (BSA, Fisher Bioreagents) and 5% normal goat serum (NGS, Gibco #PCN5000) in PBS for 1 hour at room temperature. Primary antibodies (listed in supplementary information **Table S2**) in blocking buffer were incubated overnight at 4°C followed by washing and incubation with Alexa Fluor conjugated secondary antibodies (Invitrogen, 1:1000 dilution) at room temperature for 1-2 hour in the dark. Slides were mounted with ProLong Gold Antifade reagent with DAPI (Invitrogen #P36931). Images were obtained using Leica Software on a Leica SPE confocal microscope and analyzed using ImageJ software.

### Statistical analysis

All *in vitro* experiments were repeated at least three times unless indicated otherwise. For CAM tumor growth analysis, a minimum of 5 tumors were analyzed per group/experiment. For mouse experiments, a minimum of 15 mice per group were used for tumor growth, while a minimum of 5 mice per group were used for immunostaining analysis. Statistical analysis was performed on Prism software using unpaired t-test, Mann-Whitney test, and 2-way ANOVA with Holm Sidak’s multiple comparison test. A p-value ≤ 0.05 was considered significant.

