## Supplementary Information for "Epigenetic reprogramming of DCCs into dormancy suppresses metastasis *via* restored TGFβ–SMAD4 signaling"

#### FIGURE LEGEND

**Figure S1. Supplementary data for Figure 1. a.** Principal component analysis (PCA) was generated using the top 500 genes with the highest variation in gene expression across samples. **b.** RNA expression measured by RT-qPCR in T-HEp3 cells reprogrammed with AZA+atRA (AA). **c.** Gene set enrichment analysis (GSEA) of DEGs in AZA+atRA reprogrammed T-HEp3 cells (correspond to Figure 1b). Significantly (adjusted p-value ≤ 0.05) enriched pathways are shown in the table. **d.** Top upregulated and downregulated pathways from the Wikipathways database in AZA+atRA reprogrammed T-HEp3 cells (correspond to Figure 1b). Significantly (adjusted p-value ≤ 0.05) enriched pathways are shown in the table.

**Figure S2. Supplementary data for Figure 1. a.** Graph shows the result of the chicken chorioallantoic membrane (CAM) assay. D-HEp3 cells were depleted of NR2F1 by specific siRNAs *in-vitro*. 0.5 x $10^6$ cells were inoculated to chicken CAM and grown for 7 days. After a week, tumor nodules were extracted, enzymatically digested, and counted for tumor cells. Depletion of NR2F1 in D-HEp3 cells lead to increased cell proliferation *in-vivo*. siRNA1 and siRNA2 were used for RNA sequencing experiment. **b.** Heatmap showing the genes depending on NR2F1 for AZA+atRA reprogramming. **c.** GSEA of AZA+atRA reprogrammed DEGs dependent on NR2F1. **d.** GSEA of DEGs in AZA+atRA reprogramming independent of NR2F1 (correspond to Figure 1e). Significantly enriched pathways are shown in the table.

**Figure S3. Supplementary data for Figure 2. a.** GSEA of DEGs in D-HEp3 and T-HEp3 (correspond to Figure 2a). Significantly (adjusted p-value ≤ 0.05) enriched pathways are shown in the table. **b.** ChEA of DEGs in D-HEp3 and T-HEp3 using Enrichr to compute over-representation of transcription factor targets showing top 10 hits. **c.** The Venn diagram compares upregulated and downregulated genes in D-HEp3 with upregulated and downregulated genes upon AZA+atRA reprogramming. **d.** Heatmap shows genes upregulated in T-HEp3 and regulated by enhancers. **e.** Common genes upregulated in AZA+atRA reprogramming and also regulated by enhancers in D-HEp3.

**Figure S4. Supplementary data for Figure 3. a-b.** Graphs show the change in RNA expression of TGFβ1/2/3 and TGFβ receptor 1/2/3 upon AZA+atRA reprogramming measured by RT-qPCR. **c.** Western blot shows the upregulation of SMAD4 and p27 expression upon AZA+atRA and AZA+AM80 treatment in T-HEp3 cells. Graph shows the quantification of western blot signals. **d.** Representative images of IF staining of p27 in T-HEp3 cells reprogrammed with AZA (5 nM) + AM80 (50nM). **e.** Quantification of nuclear-localized p27 (correspond to Figure S4d). **f.** Expression level of SMAD4 mRNA quantified by qRT-PCR.

**Figure S5.** Kaplan-Meier plot shows the regression free survival of HNSCC patients (n=500). **a.** SMAD2/3 (top 100 genes from NR2F1-ind)) and **b.** SMAD4 target genes (all NR2F1-ind) were used for the analysis.

**Table 1: siRNAs**

| **siRNA** | **Manufacture** | **Catalogue** |
|---|---|---|
| NR2F1 siRNA1 | Thermo Fisher | 4390824 |
| NR2F1 siRNA2 | Thermo Fisher | 4392420 |
| NR2F1 siRNA3 | Millipore Sigma | SASI_Hs01_00095428 |
| NR2F1 siRNA4 | Millipore Sigma | SASI_Hs01_00095429 |
| SMAD4 | Horizon | J-003902-09-0005 |

**Table 2: Antibodies**

| **Antibody** | **Manufacture** | **Catalogue** |
|---|---|---|
| SMAD2 | Cell Signaling Technology (CST) | 5339S |
| SMAD3 | CST | 9523P |

| | | |
|---|---|---|
| SMAD4 | CST | 38454 |
| phosphor-SMAD2 | CST | 3108S |
| Phosphor-SMAD3 | CST | 9520P |
| p27 (Western) | CST | 3686S |
| SMAD1 | CST | 9743S |
| SMAD5 | CST | 12534S |
| Phosphor-SMAD1/5 | CST | 9516S |
| ERK1/2 | BD Biosciences | 610031 |
| Phosphor-ERK1/2 | Santa Cruz Biotechnology | SC7383 |
| p38$\alpha$ | BD Biosciences | 612168 |
| Phosphor-p38 | BD Biosciences | 612288 |
| Beta-Tubulin | Sigma | T7816 |
| H3K27ac | Abcam | Ab177178 |
| Vimentin | R&D Systems | MAB2105 |
| NR2F1 | Abcam | Ab181137 |
| SMAD4 | CST | 46535S |
| p27 (IF) | CST | 3698S |
| Lamin B1 | Abcam | Ab16048 |

**Table 3: qPCR Primers**

| **Primer** | **Forward** | **Reverse** |
|---|---|---|
| NR2F1 | GCCTCAAAGCCATCGTGCTG | CCTCACGTACTCCTCCAGTG |
| TGF$\beta$1 | ATGGAGAGAGGACTGCGGAT | GAGGGAGAGAGAGGGAGTGG |
| TGF$\beta$2 | CTTTGGATGCGGCCTATT | CCCTTTGGGTTCGTGTATC |
| TGF$\beta$3 | TGGACACTTGGTTAGACGCC | GAACACAGGGTCTTGGAGGG |
| TGF$\beta$R1 | AAGTCATCACCTGGCCTTGG | ATGGTGAATGACAGTGCGGT |
| TGF$\beta$R2 | CATTTGGTTCCAAGGTGCGG | CCAGCACTCAGTCAACGTCT |
| TGF$\beta$R3 | TGAGCAGGCTGAAGTGACTG | AAACAGGAGCTCATCAGGGC |
| HOXA5 | ATGCGCAAGCTGCACATAAG | CGGGTCAGGTAACGGTTGAA |
| STRA6 | TGACAGGGACGGCCATTTAC | AGCAGGTAGGAGACATCCGT |

| | | |
|---|---|---|
| RARβ | GTACCACTATGGGGTCAGCG | CGACAGTATTGGCATCGATTCC |
| TIMP1 | AGTTTTGTGGCTCCCTGGAA | TCCGTCCACAAGCAATGAGT |
| SMAD4 | GGTAGCTGGAGAGGAAGGGA | TCAATCCAAGCCCGTGAGTC |
| TUBULIN | CCCTCCAAGCTCTACTCT | GACCAAGGCTGGTCTCTTTC |

### Figure S1

a

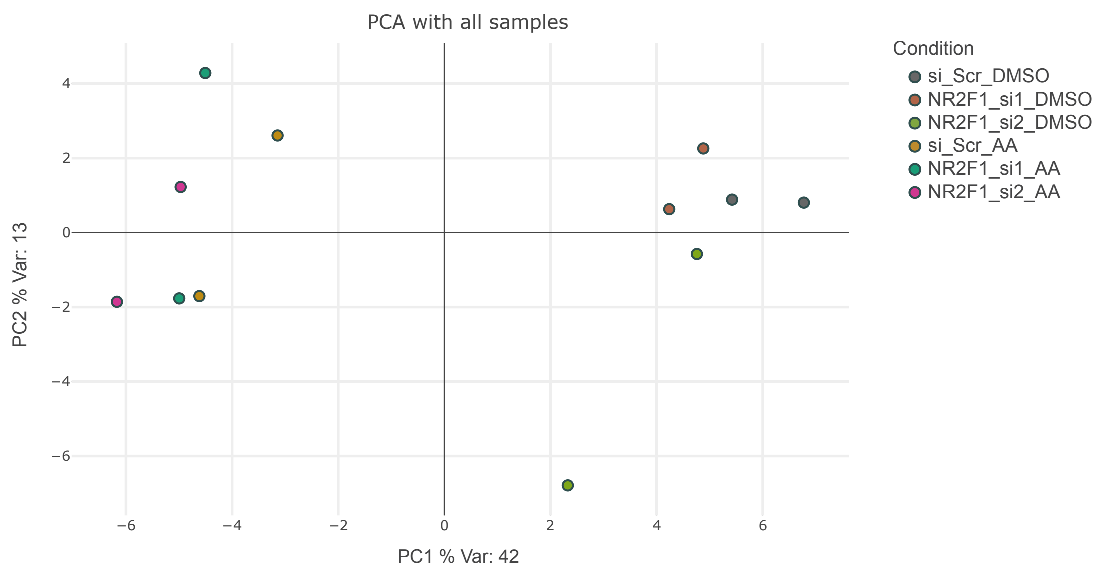

b

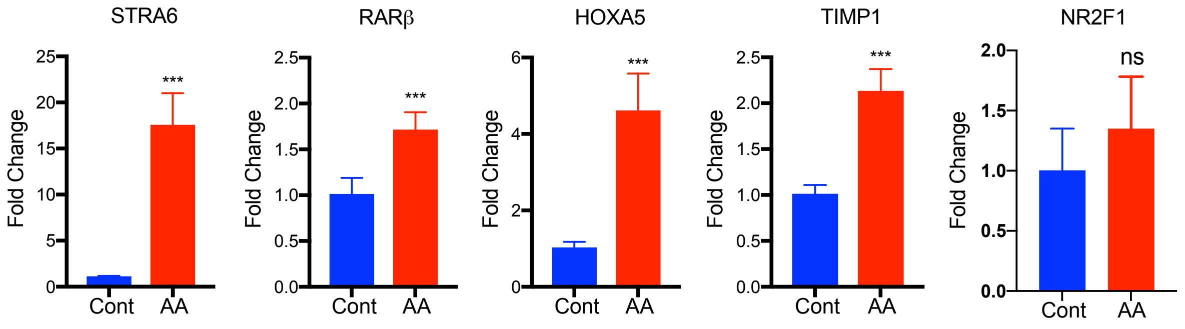

c

**Gene Set Enrichment Analysis**

| Pathway upregulated in perturbation | Adj. p-value |
|---|---|
| TNFa signaling via NFkB | 1.22E-05 |
| KRAS signaling | 0.000385978 |
| Cholesterol homeostasis | 0.000385978 |
| Estrogen response early | 0.001092615 |
| mTORC1 signaling | 0.002598046 |
| Inflammatory response | 0.02086108 |
| Androgen response | 0.03268188 |
| **Pathway downregulated in perturbation** | |
| TNFa signaling via NFkB | 1.54E-05 |
| Hypoxia | 3.60E-05 |
| UV response down | 0.001659777 |
| Androgen response | 0.029024259 |

d

**Wikipathways pathway database**

| Pathway upregulated in perturbation | Adj. p-value |
|---|---|
| Cholesterol metabolism | 1.61e-05 |
| Cholesterol biosynthesis pathway | 3.64e-04 |
| TGFbeta receptor signaling in skeletal dysplasias | 6.89e-03 |
| Notch1 regulation of human endothelial cell calcification | 2.11e-02 |
| TGFbeta receptor signaling | 2.11e-02 |
| Monoamine transport | 3.71e-02 |
| **Pathway downregulated in perturbation** | |
| Nuclear receptors metapathway | 3.15e-04 |
| Ferroptosis | 9.20e-03 |
| Phytochemical activity on NRF2 transcriptional activation | 9.20e-03 |
| NRF2 pathway | 1.06e-02 |
| Methionine de novo and salvage pathway | 1.06e-02 |
| NRF2ARE regulation | 2.04e-02 |
| Proximal tubule transport | 2.61e-02 |

### Figure S2

**a**

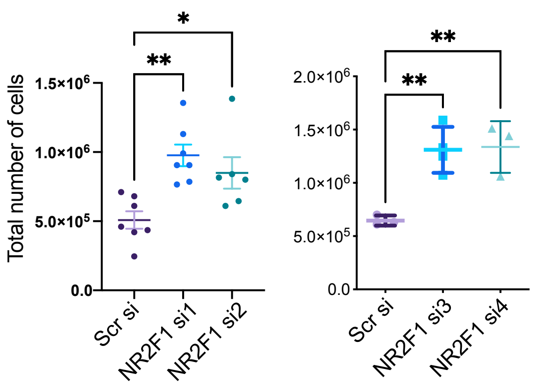

**b**

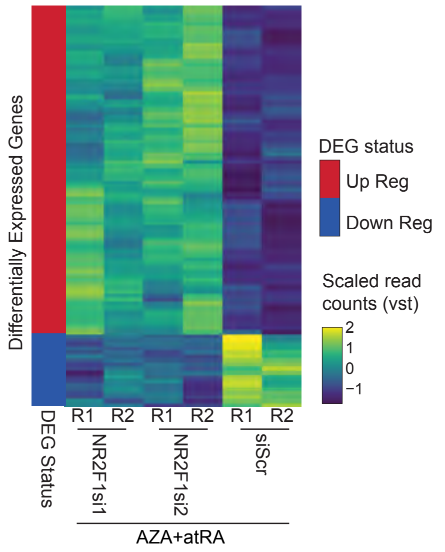

**c**

**Gene Set Enrichment Analysis**

| **Pathway upreguated in perturbation** | **Adj. p-value** |
|---|---|
| Hypoxia | 2.64e-02 |
| IL6 JAK STAT3 Signaling | 4.71e-02 |
| **Pathway downreguated in perturbation** | |
| TNFa signaling via NFkB | ns |
| KRAS Signaling dn | ns |

**d**

**Gene Set Enrichment Analysis**

| **Pathway upreguated in perturbation** | **Adj. p-value** |
|---|---|
| Cholesterol Homeostasis | 0.004140351 |
| mTORC1 signaling | 0.006653952 |
| TNFa signaling via NFkB | 0.009183079 |
| Coagulation | 0.011191592 |
| Androgen response | 0.032856885 |
| Epithelial Mesenchymal Transition | 0.037134542 |
| KRAS signaling up | 0.042799315 |
| **Pathway downreguated in perturbation** | |
| Myc target V2 | 7.12E-05 |
| Hypoxia | 0.048295865 |
| TNFa signaling via NFkB | 0.049138068 |

### Figure S3

**a**

**Gene Set Enrichment Analysis**

| Pathway upreguated in D-HEp3 | Adj. p-value |
|---|---|
| E2F Targets | 0 |
| mTORC1 signaling | 0 |
| Myc targets V1 | 0.004048583 |
| Myc targets V2 | 0.020833334 |
| G2M Checkpoint | 0.003984064 |
| Oxidative phosphorylation | 0.007604563 |
| Glycolysis | 0.014084507 |
| Fatty acid metabolism | 0.026229508 |
| **Pathway upreguated in T-HEp3** | |
| Coagulation | 0 |
| Epithelial mesenchymal transition | 0 |
| TNFA signaling via NFkB | 0 |
| KRAS signaling up | 0.00152207 |
| IL2 STAT5 signaling | 0.002881844 |
| Hedgehog signaling | 0.012367492 |
| Inflammatory response | 0.005830904 |
| Angiogenesis | 0.04676259 |
| Complement | 0.02209131 |
| Myogenesis | 0.036418818 |
| Apoptosis | 0.024745269 |
| IL6 JAK STAT3 signaling | 0.06319115 |
| Xenobiotic metabolism | 0.05950653 |
| Allograft rejection | 0.06962025 |
| Mitotic spindle | 0.05 |

**b**

**ChEA 2016**

**D-HEp3**

| Enriched TFs up genes | Adj. p-value |
|---|---|
| EKLF | 3.76E-29 |
| XRN2 | 6.83E-28 |
| GABP | 1.98E-27 |
| FOXM1 | 7.28E-24 |
| PRDM5 | 1.22E-22 |
| ETS1 | 9.56E-22 |
| MYC | 2.11E-20 |
| JARID1A | 7.93E-20 |
| E2F4 | 5.14E-19 |
| DACH1 | 2.49E-17 |

**T-HEp3**

| Enriched TFs up genes | Adj. p-value |
|---|---|
| CREM | 6.05E-40 |
| NUCKS1 | 1.01E-25 |
| MITF | 3.58E-23 |
| P300 | 7.61E-23 |
| NCOR | 1.07E-22 |
| KDM2B | 8.09E-22 |
| BRD4 | 2.98E-20 |
| KLF6 | 7.45E-20 |
| ELF3 | 1.87E-19 |
| CREB1 | 9.34E-19 |

**c**

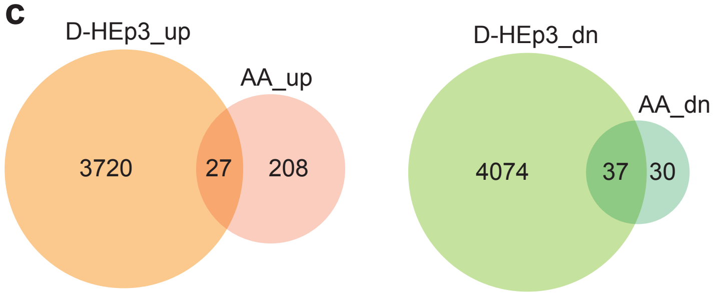

**d**

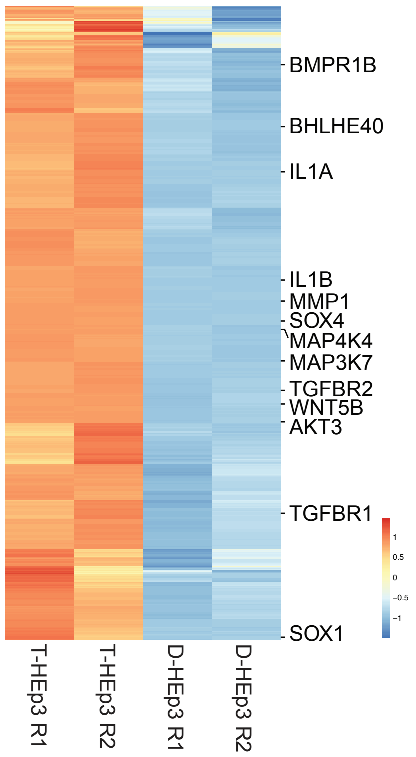

**e**

**Common (SMADs target) genes in AZA+atRA reprogramming and regulated by enhancer in D-HEp3 cell line**

| SMAD2/SMAD3 | SMAD4 |
|---|---|
| RP1L1 | KRT15 |
| KRT15 | DHRS3 |
| DHRS3 | PMEPA1 |
| LIF | LAMB3 |
| CYP26B1 | MBNL2 |
| KRT8 | |
| LAMB3 | |
| OSBPL7 | |
| MAP3K9 | |
| MBNL2 | |
| IFRD1 | |

### Figure S4

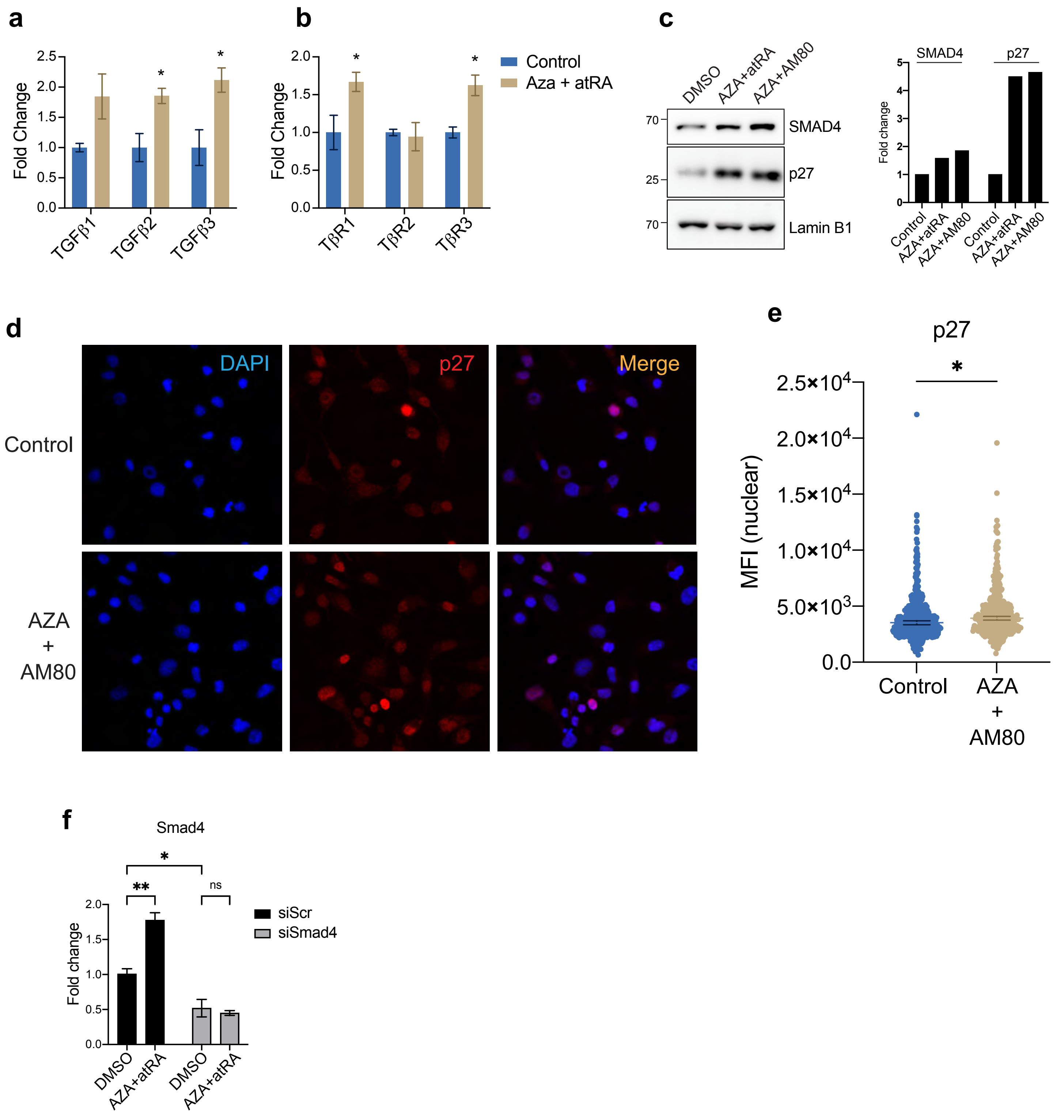

a
Fold Change
2.5
2.0
1.5
1.0
0.5
0.0
*
*
TGFβ1
TGFβ2
TGFβ3
b
Fold Change
2.0
1.5
1.0
0.5
0.0
*
*
TβR1
TβR2
TβR3
Control
Aza + atRA
c
DMSO
AZA+atRA
AZA+AM80
70
25
70
SMAD4
p27
Lamin B1
SMAD4
p27
Fold change
5
4
3
2
1
0
Control
AZA+atRA
AZA+AM80
Control
AZA+atRA
AZA+AM80
d
DAPI
p27
Merge
Control
AZA
+
AM80
e
p27
*
MFI (nuclear)
2.5×10^4
2.0×10^4
1.5×10^4
1.0×10^4
5.0×10^3
0.0
Control
AZA
+
AM80
f
Smad4
*
**
ns
Fold change
2.0
1.5
1.0
0.5
0.0
DMSO
AZA+atRA
DMSO
AZA+atRA
siScr
siSmad4

### Figure S5

a

SMAD2/SMAD3 Target genes

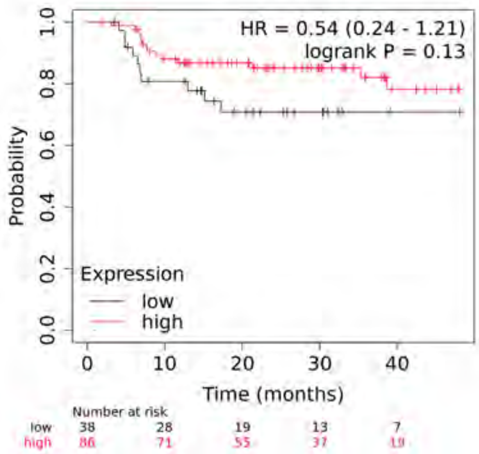

Regression Free Survival

b

SMAD4 Target genes

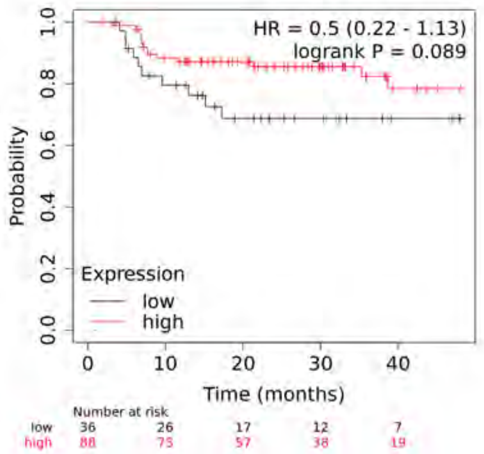

Regression Free Survival
